# Enzyme-less nanopore detection of post-translational modifications within long polypeptides

**DOI:** 10.1101/2023.02.09.527483

**Authors:** Pablo Martin-Baniandres, Wei-Hsuan Lan, Stephanie Board, Mercedes Romero-Ruiz, Sergi Garcia-Manyes, Yujia Qing, Hagan Bayley

## Abstract

Means to sequence DNA and RNA quickly and cheaply have revolutionized biology and medicine. The ability to analyse cellular proteins and their millions of variants would be an advance of comparable importance, but requires a fresh technical approach. We use electroosmosis for the non-enzymatic capture, unfolding and translocation of individual polypeptides of more than 1200 residues by a protein nanopore. By monitoring the ionic current carried by the nanopore, we locate post-translational modifications deep within the polypeptide chains, and thereby lay the groundwork for obtaining inventories of the proteoforms in cells and tissues.

---

Single-molecule nanopore proteomics is gaining momentum^1^. Nanopore sequencing of ultralong DNA and RNA has enabled applications in basic science and medicine that challenge short-read technologies^2^. Modulation of the ionic current passing through a nanopore might also be used to distinguish and count the millions of proteoforms expressed from the 20,000 or so protein-encoding human genes. In this way, inventories would be obtained of variations such as post-translational modifications (PTMs) and alternative RNA splicing, which are often present at multiple locations throughout a polypeptide chain^3^. While recent studies with nanopores have mainly examined short peptides^4^,^5^, knowledge of the architecture of long polypeptide chains would be far more informative, but obtaining such information encounters two main roadblocks. First, proteins adopt tertiary structures that prohibit nanopore translocation. Second, unlike DNA or RNA, polypeptides have a low-density and heterogenous distribution of charge along their chains, which renders electrophoresis inapplicable as a means of translocation. One solution is to incorporate charged leader sequences, such as a single-stranded DNA^6–8^, to implement the electrophoretic capture and unfolding of proteins. However, as soon as the leader sequence exits the pore, a directional force is no longer present, and if the remaining polypeptide unfolds, it will diffuse in a partially extended conformation through the pore^6–8^. In another approach, unfoldases (e.g., ClpX^9^ or VATΔN^10^) have been employed to drive the translocation of tagged polypeptides through nanopores after electrophoretic threading. Unlike the ratcheting enzymes used for DNA and RNA nanopore sequencing^11–13^, these unfoldases are not capable of residue-by-residue translocation of polypeptides. Further, whether large PTMs will be tolerated during enzymatic translocation is unclear. Therefore, we aimed to establish a general non-enzymatic means to map modifications within full-length polypeptide chains, and eventually to inventory the collection of proteoforms in individual cells, rather than perform an ensemble analysis of peptide fragments.

Previously, we distinguished C-terminal PTMs (phosphorylated serines) in a model protein by using the DNA-leader approach^7^. However, the detection by nanopores of PTMs deep within a long polypeptide chain has remained a challenge. As an alternative to electrophoresis, electroosmotic flow (EOF) within a charge-selective protein nanopore has been shown to modulate the binding and dissociation of neutral small molecules^14^, and promote the trapping of short peptides or folded proteins^15–17^. Recently, simulations have suggested an electroosmotic contribution to the translocation of polypeptides through the wild-type (WT) staphylococcal α-hemolysin (αHL) following electrophoretically assisted threading^18^. The ionic charge selectivity necessary to generate electroosmotic flow was attributed, computationally, to guanidinium cations lining the interior of the pore^18^. Here, we use strong electroosmosis directly attributable to the charged side chains in an engineered αHL pore to capture long underivatized polypeptides and detect modifications within them as they are unfolded and translocated.

## Electroosmotic translocation of protein concatemers

We constructed dimers, tetramers, hexamers and octamers of thioredoxin (**Fig. 1a-b, Supplementary Table 1, Fig. 1**). The thioredoxin (Trx, 108 amino acids) had the two catalytic cysteines removed (Trx: C32S/C35S)^6^. The Trx monomers were connected by 29-amino acid linkers, capable of spanning the 10-nm long lumen of the αHL nanopore when fully extended (0.35 nm per aa). An exception was that the Trx-linker octamer had no C-terminal linker (Supplementary Table 1). We used an anion-selective αHL mutant, (NN_113R)_7_ (P_Na+_/P_Cl-_ = 0.33^14^), to generate electroosmosis. All four Trx-linker concatemers were captured by the (NN_113R)_7_ in the presence of 750 mM guanidinium chloride (GdnHCl) (**Fig. 1c**) at a capture rate ~25 times faster than that of a WT αHL pore (P_Na+_/P_Cl-_ = 0.78^14^) (*k*_(octamer)_ ~2.5 s^-1^μM^-1^ with (NN_113R)_7_ versus ~0.11 s^-1^μM^-1^ with (WT)7, recording conditions: 750 mM GdnHCl, 10 mM HEPES, pH 7.2, +140 mV (trans), 24 ± 1 °C).

**Fig. 1.**
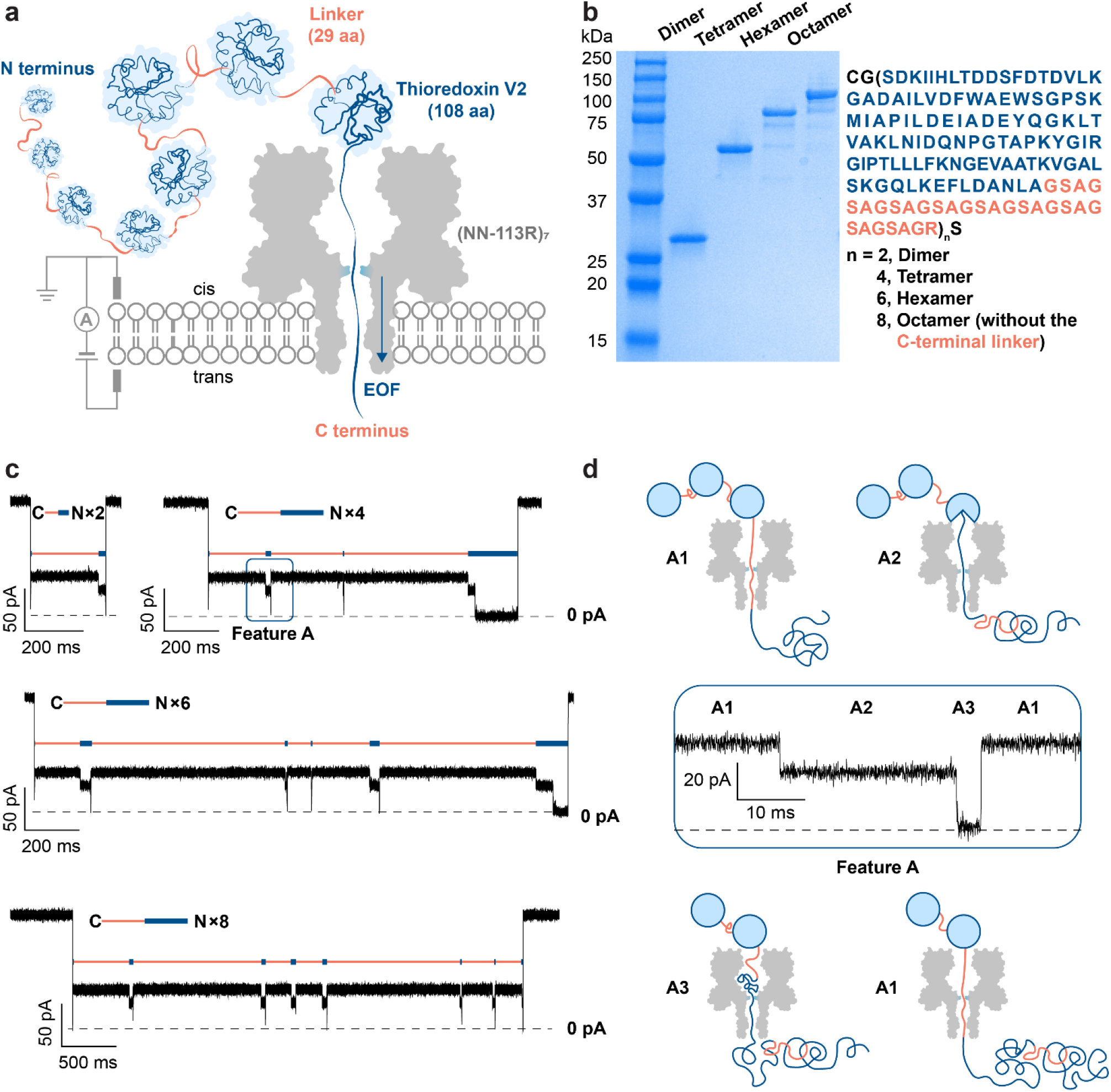
Electroosmosis-driven translocation of thioredoxin-linker concatemers through a protein nanopore. **a,** Electroosmotic flow (EOF) in a charge-selective α-hemolysin nanopore, (NN-113R)_7_, drives the sequential co-translocational unfolding of thioredoxin (Trx) units within a polyprotein of >1000 amino acids. **b**, An SDS-polyacrylamide gel showing the Trx-linker dimer (28 kDa), tetramer (55 kDa), hexamer (83 kDa), and octamer (110 kDa). **c**, Current recordings for the C terminus-first translocation of a dimer, a tetramer, a hexamer and an octamer without post-acquisition filtering. The repeating features A are indicated by orange and blue bars. **d**. Zoom-in of the repeating feature A boxed in blue in panel **c** without post-acquisition filtering. Three levels are assigned as: A1. a linker within the pore; A2, A3. different segments of partly unfolded Trx within the pore. Conditions in **c** and **d**: 750 mM GdnHCl, 10 mM HEPES, 5 mM TCEP, pH 7.2, Trx-linker concatemers (cis) (dimer: 2.23 μM; tetramer: 0.63 μM; hexamer: 0.25 μM; octamer: 0.81 μM), +140 mV (trans), 24 ± 1 °C.

Electroosmosis-driven concatemer translocation produced current patterns containing repeating features (**Fig. 1c, Supplementary Fig. 2**). The most abundant feature, A, consisted of three levels (A1, A2, A3) (**Fig. 1c-d**). The percentage residual current (I_res%_) for each level in feature A was consistent across all such events for each polypeptide translocation and between all individual concatemers observed with the same or different pores (**Supplementary Table 2**). A spike to ~0 pA was seen at the beginning of almost all the translocation events and was speculated to represent the rapid unfolding and translocation of the first Trx-linker unit. The spike was followed by up to n-1 repeats of the three-step feature A (n = number of Trx-linker units in the concatemer), which unambiguously demonstrated the stepwise translocation of entire polypeptide chains one unit at a time.

Less often, a different repeating element, B, was recorded (**Supplementary Fig. 2a, Table 3**). Further, when two identical concatemers were linked by a disulfide bond between the N-terminal cysteines, feature B occurred only after feature A within each translocation event (**Supplementary Fig. 1b**). Therefore, we assigned these two features as C terminus-first (A) and N terminus-first (B) translocation events. In the presence of a C-terminal linker (e.g. Trx tetramer, **Fig. 1b**), the ratio of C terminus-versus N terminus-first translocation events was ~2:1 at +140 mV (**Supplementary Table 3**); in the absence of the C-terminal leader sequence (e.g. Trx octamer, **Fig. 1b**), the C terminus-first pattern dominated (C versus N = ~10:1 at +140 mV) (**Supplementary Table 3**).

The repeating feature A was lost at a GdnHCl concentration of 3 M (**Supplementary Fig. 3**). At 750 mM GdnHCl, ~12% of the translocated octamers produced the maximum of 7 repeats of feature A following the initial spike; kinetic analysis revealed two populations of A3: one had a mean dwell time ~500 times longer than the other at +140 mV (<T_A3_> = 320 ± 60 ms versus 0.69 ± 0.04 ms) (**Supplementary Table 4**). The longer-lived A3 (τ_A3_>10 ms) was seen in 25% of the final features A recorded as translocation of an octamer was completed, but only in 3% of the preceding features A. Tentatively, we assign Level A1 as a threaded linker preceding the C-terminus of a folded Trx unit; Level A2 as a C-terminal portion of a partially unfolded Trx unit extended into the nanopore; Level A3 as the spontaneous unfolding and passage of the remaining Trx polypeptide through the nanopore (**Fig. 1d**). The dominant absence of a multi-level feature for the first unit and an extended duration for the last unit suggest that the unfolding kinetics of Trx units differ when the polypeptide chain is unable to fully span the lumen of the nanopore.

Similar repeating features were also seen in the absence of GdnHCl (recording conditions: 750 mM KCl, 10 mM HEPES, pH 7.2, +140 mV (trans), 24 ± 1 °C) or in the presence of a non-denaturing concentration of urea (recording conditions: 2 M urea, 750 mM KCl, 10 mM HEPES, pH 7.2, +140 mV (trans), 24 ± 1 °C). Under these conditions, the translocation kinetics were slower (**Fig. 2**) and these options were not pursued further.

**Fig. 2.**
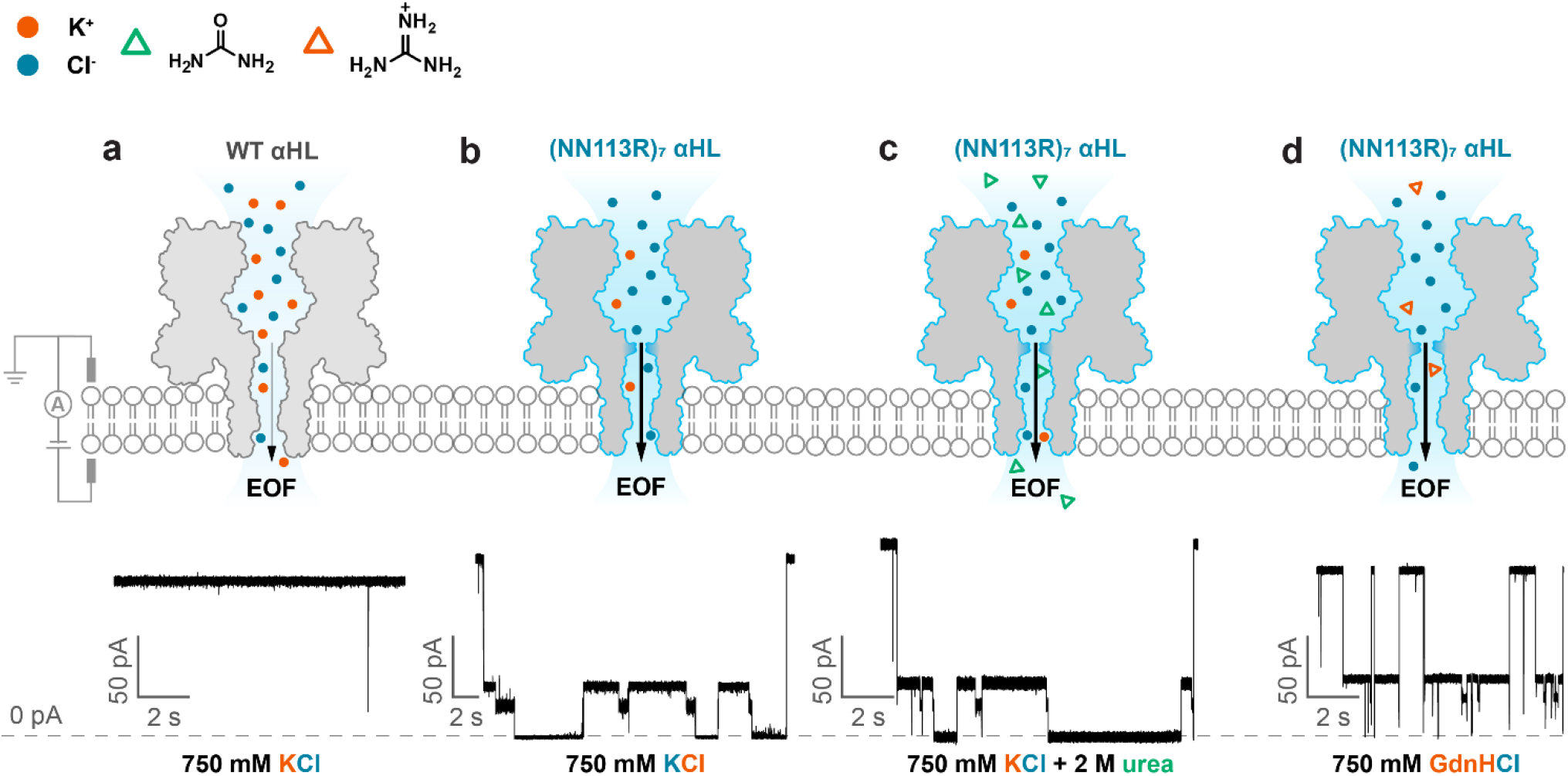
Chaotrope-facilitated electroosmotic translocation of the Trx-linker octamers through a nanopore. **a,** The translocation of Trx-linker octamers through a weakly anion-selective wild-type (WT) α-hemolysin (αHL) was not observed in the absence of a chaotrope. **b-d**, Current traces showing the translocation of Trx-linker octamers through the electroosmotically active nanopore, (NN-113R)7, in the presence of 750 mM KCl **(b)**, 750 mM KCl and 2 M urea **(c)**, or 750 mM GdnHCl **(d)**with 2 kHz post-acquisition filtering. The use of non-denaturing concentrations of chaotropic agents (urea and guanidinium chloride, GdnHCl) accelerated the co-translocational unfolding of the Trx units. Conditions: 10 mM HEPES, pH 7.2, 0.81 μM Trx-linker octamer (cis), +140 mV (trans), 24 ± 1 °C, with **(a-b)**750 mM KCl; **(c)**2 M urea and 750 mM KCl; **(d)**750 mM GdnHCl.

## Detection of post-translational modifications during electroosmotic translocation

To determine whether PTMs near the middle of a long polypeptide chain could be located during electroosmosis-driven translocation, we constructed Trx-linker nonamers containing a modification site (RRASAC) at two different positions in the central linker (**Supplementary Table 1, Fig. 4**) for serine phosphorylation (14S-P or 24S-P) or cysteine-directed glutathionylation or glycosylation (16C-GSH, 26C-GSH, 16C-SLN, or 26C-SLN) (**Fig. 3a**). In the presence of a phosphate group (P) or glutathione (GSH) or 6’-sialyllactosamine (SLN), Level A1 for the modified units exhibited a smaller I_res%_ and higher root-mean-square noise (I_RMS_) than that of unmodified segments within an individual polypeptide (**Fig. 3b, Supplementary Table 5**). The average increment in the current blockade was roughly proportional to the mass of the PTM with phosphate giving the smallest increment and the trisaccharide the largest (**Supplementary Table 5**), although there was substantial overlap between the 14S-P/24S-P and 16C-GSH/26C-GSH populations (**Fig. 3b, Supplementary Fig. 5**). Mixed nonamers containing different PTMs at the same site (26C-GSH and 26C-SLN) were discriminated using a single pore (**Supplementary Fig. 6**). All three PTMs tested caused a smaller current blockade at serine 14 (14S) or cysteine 16 (16C) than at serine 24 (24S) or cysteine 26 (26C) (**Fig. 3b**). Given that 14S/16C must be closer to the cis opening of the αHL pore than 24S/26C in a C terminus-first threading configuration, it is likely that the central constriction of the pore is located closer to 24S/26C (**Fig. 3a, Supplementary Fig. 7**). The findings also suggest that the polypeptide might not be fully extended under the EOF (**See Supplementary Fig. 7 for further analysis**), which corroborates force spectroscopy data for polypeptides under forces at <20 pN^19^,^20^.

**Fig. 3.**
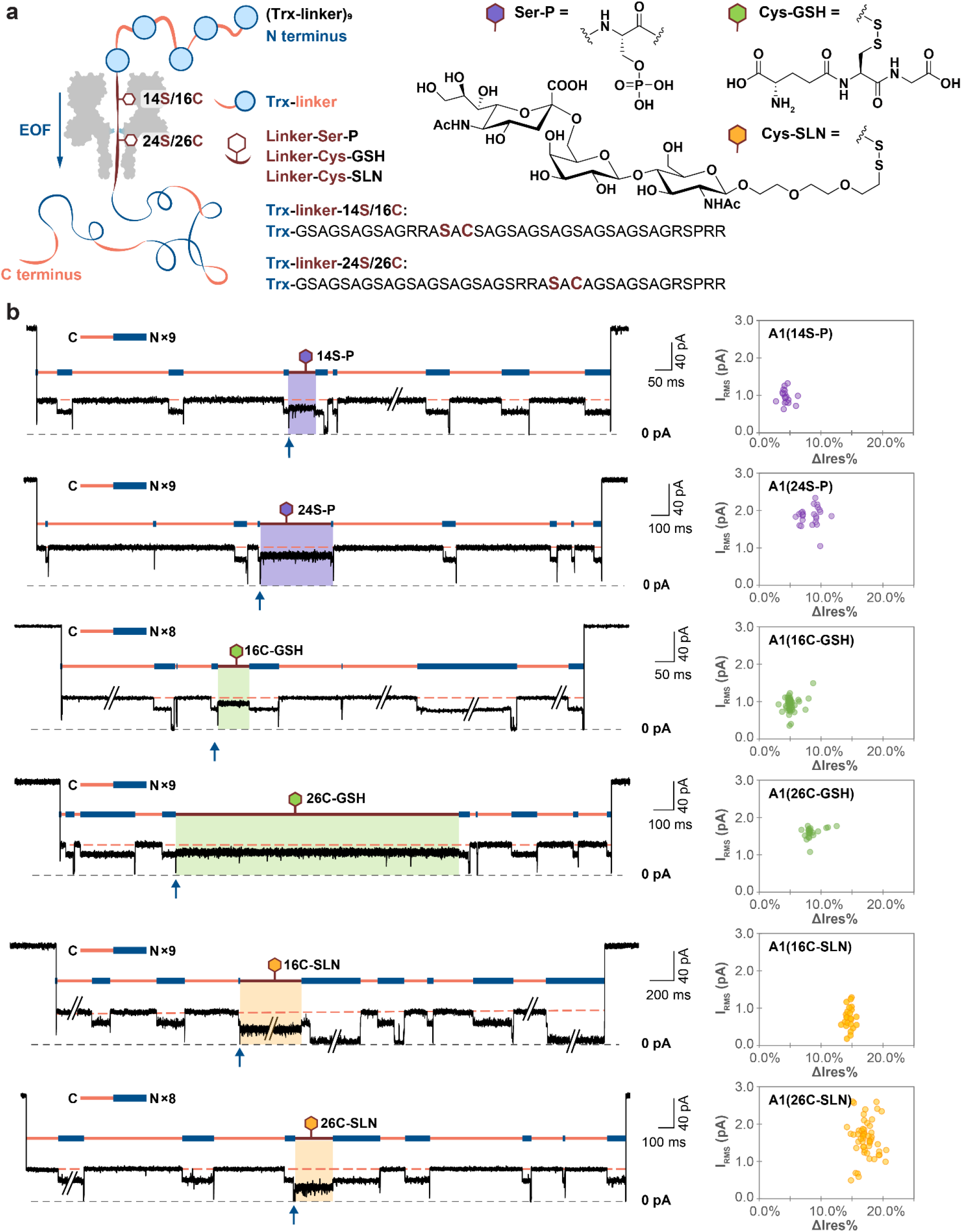
Detection of PTMs in protein concatemers traversing a nanopore driven by electroosmotic flow. **a,** The Trx-linker nonamers tested contained a RRASAC sequence within the central linker, which was post-translationally phosphorylated (purple), S-glutathionylated (green) or glycosylated (yellow). **b,** Left: Recordings of C terminus-first translocation events of Trx-linker nonamers showing a distinct Level A1 (boxed in purple, green or pink) in the presence of a PTM compared to the unmodified A1 (orange dash). Traces have been filtered at 2 kHz; transient A3 levels were truncated by filtering and therefore deviated from ~0 pA. The A3 produced by the translocation of an unmodified unit before the modified linker is indicated with a blue arrow and each of the features A recorded is indicated by orange and blue bars. The number of repeats of feature A is specified. Right: Scatter plots of I_RMS_ and ΔI_res%_ for individual translocation events, ΔI_res%_ = <I_res%_(A1, Trx-linker)> – I_res%_(A1, Trx-linker+PTM), where <I_res%_(A1, Trx-linker)> is the mean I_res%_ value of the remaining A1 levels for unmodified repeat units within an individual translocation event. Conditions: 375 mM GdnHCl, 375 mM KCl, 10 mM HEPES, pH 7.2, 1.2 μM Trx-linker nonamer (cis), +140 mV (trans), 24 ± 1 °C.

## Conclusions

Here, we have established that electroosmotically active nanopores can capture and unfold individual proteins comprising long (>1200 aa) polypeptide chains for PTM identification and localisation. To a first approximation, the electroosmotic force acting on a polypeptide remains constant during translocation, which creates a unidirectional bias desirable for the placement of PTMs in sequence. In contrast, the overall time for unforced polypeptide translocation scales roughly as the square of its length, because the polypeptide chain can move back and forth before diffusing out of the pore^21^. This is the case within a electroosmotically-inactive nanopore after the exit of a charged leader sequence^6^ or immediately after a protein domain has unfolded during movement propelled by a motor protein^9^, which is not ideal for the sequential detection of modification sites within individual polypeptide chains. As a label-free method, our approach circumvents the need to derivatize proteins at either the N or C terminus for electrophoretic translocation, which could be problematic for eukaryotic proteins due to the widespread presence of N-acetylation and the lack of efficient N or C terminus-specific chemistries.

Here, we have located PTMs in linkers deep within long polyprotein chains by exploiting stepwise unfolding. Our encouraging results lay the groundwork for building inventories of underivatised full-length proteoforms from cells and tissues. The detection of PTMs in freely translocating domains will require the slowing and stretching of polypeptide chains, which might be produced by physical effects (heat, voltage ramps etc), nanopores with different internal geometries and surface charge, and the use of weakly bound ligands. Recording at MHz acquisition rates^22^ might also prove advantageous. These endeavours are beyond the scope of the present work and indeed depend on the advances we report here.

Our strategy will be readily transferable to nanopore sequencing devices (e.g., the MinION) for highly parallel PTM profiling, which will be useful for producing inventories of full-length human proteoforms, which are ~500 aa in median length^23^. To achieve the complete characterisation of the proteoforms in individual cells, our approach is faced with challenges. For example, because proteins vary in their resistance to unfolding, it will be problematic to establish universal conditions for the translocation of all the protein components of a cell. Similarly, the capture rate is likely to differ between proteins. To this end, voltage sweeps might be used in combination with denaturants to promote protein capture and enable co-translocational unfolding. Further, compact PTMs (e.g. methylation) might be challenging to detect directly. Ligand-assisted detection might be attempted by the use of antibodies or chemical binders. In summary, our enzyme-less approach, targeting full-length proteins, presents a viable nanopore technology, which will ultimately allow comprehensive proteoform inventories to be established for tissues and single cells. These massive sets of information will extend beyond what is recognised from DNA and RNA sequencing, and will potentially unveil as yet unknown aspects of the biology of cells and tissues.

## Methods

### Construction of Trx-linker concatemer genes

All reagents were purchased from NEB (New England Biolabs) and DNA oligonucleotides were obtained from IDT (Integrated DNA Technologies) unless otherwise indicated. Trx-linker concatemer genes were prepared as previously described^24^. Briefly, the Trx-linker monomer gene was amplified with a 5’ primer containing a BamHI restriction site and a 3’ primer containing a BglII restriction site, which permitted in-frame cloning of the monomer into the vector pQE30 (Qiagen). Synthetic genes encoding the concatemers were then constructed by iterative cloning of monomer into monomer, dimer into dimer, and tetramer into tetramer. To aid purification, an N-terminal SUMO tag was inserted between the His6 tag and the first monomer unit. In addition, N-terminal cysteine-glycine codons were included to give the final concatemer constructs: His6-SUMO-CysGly-(Trx-linker)_n_ (n = 2, 4, 6) and His6-SUMO-CysGly-(Trx-linker)_7_Trx.

To produce Trx-linker nonamers (His6-SUMO-(Trx-linker)_n_, n =9) containing a modification site, the N-terminal cysteine-glycine codons were removed from the tetramer gene and a DNA cassette was designed to contain two terminal restriction sites (BamHI and BglII) and two internal restriction sites (KpnI and AvrII) (5’-p GATCCGGTACCGGCGGTCCTAGGAGATCTGGCGGTA-3’, 5’-p GCCATGGCCGCCAGGATCCTCTAGACCGCCATTCGA-3’). Using the interactive cloning strategy described above, a “cloneable” Trx-linker octamer gene was assembled with the DNA cassette as the middle unit flanked by two Trx-linker tetramer genes (i.e., the final construct is His6-SUMO-(Trx-linker)_4_-KpnI-AvrII-(Trx-linker)_4_). A Trx-linker monomer mutant gene encoding a RRASAC peptide motif was created by site-directed insertion (Forward primer: 5’-AGCGCCTGCGCGGgTtCTGCTGGTTCC-3’; Reverse primer: 5’-CGCaCgGCG GCTCCCTGCACTTCCGGC-3’) and subsequently cloned in between the KpnI and AvrII sites within the Trx-linker octamer to give (Trx-linker)_4_-Trx-linker(RRASAC)-(Trx-linker)_4_. The placement of a single correctly oriented insert was confirmed by sequencing using primers targeting the KpnI and AvrII ligation sites (Forward primer: 5’-TGCGAGCGCCTGCGGTGG-3’; Reverse primer: 5’-ACGCTCGCGGACGCCACC-3’).

### Expression and purification of Trx-linker concatemers

Genes encoding the N-terminal His6-SUMO tagged concatemers of Trx were cloned into the pOP3SU plasmid (kindly provided by Marko Hyvönen). BLR(DE3) competent cells (Novagen) were transformed with the plasmids and grown in Luria broth (LB) medium supplemented with ampicillin (100 μg/mL) at 37 °C with continuous shaking (250 rpm). Protein expression was induced in the exponential growth phase (OD_600_ = 0.6) with isopropyl-β-D-1-thiogalactopyranoside (IPTG) (0.5 mM final concentration). After 8 h, cells were harvested by centrifugation (10 min, 5,000 g), resuspended in binding buffer (30 mM Tris HCl, 250 mM NaCl, 25 mM imidazole, pH 7.2) supplemented with a protease inhibitor cocktail (cOmplete^™^, EDTA-free, Roche) and lysed by sonication. Cell debris was removed by centrifugation at 20,000 g for 45 min, and the supernatant loaded onto a HisTrap HP column (5 mL, Cytiva) at 0.2 mL/min. The column was washed with 50 mL of the binding buffer before a single step elution with 30 mM Tris HCl, 250 mM NaCl, 300 mM imidazole, pH 7.2. A single peak containing the almost pure protein was collected and dialysed (Slide-A-Lyzer G2 Dialysis Cassette, 10,000 MWCO 30 mL, ThermoFisher) for 3 h against 4 L of dialysis buffer (50 mM Tris HCl, 250 mM NaCl, 2 mM 1,4-dithio-D-threitol (DTT), pH 8.0), at 4 °C with continuous stirring, to remove excess imidazole. After injecting the His6-tagged Ulp1 protease into the dialysis cassette at a molar concentration ratio of 1:200 (Ulp1:Trx-linker concatemer), the mixture was transferred into fresh dialysis buffer overnight for SUMO-tag cleavage. The cassette was then transferred one last time into fresh dialysis buffer without DTT for 4 h. The dialysed protein was loaded onto a column packed with HisPur Ni-NTA Agarose Resin (5 mL, ThermoFisher) equilibrated with binding buffer (50 mM Tris HCl, 250 mM NaCl, pH 8.0) and the flow through was re-applied 5 more times. The final flow through containing the His6-SUMO-free protein was aliquoted and flash frozen for storage at −80 °C.

### Expression and purification of SUMO protease Ulp1

The Pfget19_Ulp1 plasmid (Addgene) containing a His6-tagged Ulp1 gene was transformed into T7 Express competent cells (NEB) and grown in LB medium supplemented with ampicillin (100 μg/mL) at 37 °C with shaking (250 rpm). Expression was induced at OD_600_ = 0.5 with IPTG (0.5 mM). Cells were harvested after 3 h by centrifugation, resuspended in lysis buffer (4 mL/ g: 50 mM Tris HCl, 300 mM NaCl, 10 mM imidazole, pH 7.5) supplemented with lysozyme (1 mg/mL), and incubated on ice for 30 min before sonication. The lysate was spun at 20,000 rpm for 45 min to remove cell debris and the supernatant was applied to a column packed with HisPur Ni-NTA Agarose Resin (5 mL, ThermoFisher) and equilibrated with binding buffer (50 mM Tris HCl, 300 mM NaCl, pH 7.5). The column was washed with 10 column volumes of wash buffer (50 mM Tris HCl, 300 mM NaCl, 20 mM imidazole, pH 7.5) and the protein was eluted with 10 mL of elution buffer (50 mM Tris HCl, 300 mM NaCl, 300 mM imidazole, pH 7.5). The eluted protein was dialysed against storage buffer (50 mM Tris HCl, 200 mM NaCl, 2 mM 2-mercaptoethanol) overnight, aliquoted and flash frozen as a 50% stock in glycerol.

### Phosphorylation of Trx-linker concatemers

Trx-linker concatemers (1 mg/mL) were incubated with 50,000 units of the catalytic subunit of cAMP-dependent protein kinase (PKA) (NEB)—which recognizes the RRAS motif within the central linker of the Trx-linker nonamer—in protein kinase buffer (50 mM Tris HCl, pH 7.5,10 mM MgCl2, 0.1 mM EDTA, 4 mM DTT, 0.01% Brij 35, and 2 mM ATP) (NEB) at 30 °C for 1 h. The solution was then supplemented with additional 2 mM ATP and 2 mM DTT before overnight incubation at 30°C. Trx-linker concatemers were purified and concentrated using centrifugal filters (Amicon Ultra-0.5 mL 100K), aliquoted and flash frozen for storage at −20°C (10 mM HEPES, pH 7.2, and 750 mM KCl). Phosphorylation of the Trx-linker concatemers at a single site was verified by LC-MS.

### Modification of cysteines on Trx-linker concatemers

Reagents were purchased from Sigma-Aldrich unless otherwise indicated. Trx-linker nonamer was first treated with tris(2-carboxyethyl)phosphine (TCEP) (70 to 100 eq) at 32 °C for 2 h in protein storage buffer (50 mM Tris HCl, 250 mM NaCl, pH 8.0). Excess TCEP was removed by a desalting column (PD MiniTrap G-25 column, Cytiva). To glutathionylate the Trx-linker nonamer, the reduced protein was reacted with oxidized glutathione (100 eq) at 32 °C overnight in protein storage buffer (50 mM Tris HCl, 250 mM NaCl, pH 8.0) before desalting to remove the excess reagent. The modified proteins were aliquoted and flash frozen for storage at −20°C. To glycosylate the Trx-linker nonamers, reduced protein was reacted first with 2,2’-dithiodipyridine (DPS) (20 eq) at 32 °C overnight in the protein storage buffer (50 mM Tris HCl, 250 mM NaCl, pH 8.0). After removal of excess DPS with a desalting column, the activated nonamer was reacted with the 6’-sialyllactosamine derivative (NeuAcα(2-6)LacNAc-PEG3-Thiol, 5 eq, Sussex Research Laboratories) overnight at 32 °C in protein storage buffer (50 mM Tris HCl, 250 mM NaCl, pH 8.0). Modified nonamers were desalted (PD MiniTrap G-25 column, Cytiva), aliquoted and flash frozen for storage at −20°C. That glutathionylation or glycosylation occurred at single sites was verified by LC-MS.

### Single-channel recording

Planar lipid bilayers of 1,2-diphytanoyl-sn-glycero-3-phosphocholine (Avanti Polar Lipids) were formed by using the Müller-Montal method on a 50 μm-diameter aperture made in a Teflon film (25 μm thick, Goodfellow) separating two 500 μL compartments (cis and trans) of the recording chamber. Each compartment was filled with recording buffer (750 mM GdnHCl, 1.5 M GdnHCl, 3 M GdnHCl, 2 M urea/750 mM KCl, or 750 mM KCl, 10 mM HEPES, 5 mM TCEP, pH 7.2 for Trx-linker dimer, tetramer, hexamer, and octamer; 375 mM GdnHCl/375 mM KCl, 10 mM HEPES, pH 7.2 for Trx-linker nonamers). To record with Trx-linker dimer, tetramer, hexamer, or octamer and ensure a reduced N-terminal cysteine, pre-treatment of the protein samples with 5 mM TCEP was carried out for 10 min at room temperature. Trx-linker concatemers were added to the cis compartment (dimer: 2.2 μM; tetramer: 0.63 μM; hexamer: 0.25 μM; octamer: 0.81 μM; nonamer: 1.2 μM). Ionic currents were measured at 24 ± 1 °C by using Ag/AgCl electrodes connected to the headstage of an Axopatch 200B amplifier. After a single (NN-113R)_7_ pore had inserted into the bilayer, the solution was replaced with fresh buffer by manual pipetting, to prevent further insertions. Signals were low-pass filtered at 10 kHz and sampled at 50 kHz with a Digidata 1440A digitizer (Molecular Devices). Current traces were idealized by using Clampfit 10.3 (Molecular Devices). The idealized data were analyzed with QuB 2.0 software (https://qub.mandelics.com/). Dwell time analysis was performed by using the maximum interval likelihood algorithm of QuB.

## Acknowledgements

This research was supported by the Wellcome Leap Delta Tissue Program, Oxford Nanopore Technologies and a European Research Council Advanced Grant (SYNTISU). The work was supported in part by the Francis Crick Institute which receives its core funding from Cancer Research U.K. (FC001002), the U.K. Medical Research Council (FC001002), and the Wellcome Trust (FC001002). W.-H.L. is funded by an Oxford-Taiwan Graduate Studentship in partnership with a Department of Chemistry Scholarship, University of Oxford. S.G.M. is supported by a Leverhulme Trust Research Leadership Award (RL 2016-015), a Wellcome Trust Investigator Award (212218/Z/18/Z) and a Royal Society Wolfson Fellowship (RSWF/R3/183006). Y.Q. was supported by a Glasstone Research Fellowship and a Fellowship by Examination, Magdalen College, Oxford.

## Author contributions

P.M.B., W.-H.L., M.R.R, S.G.-M., Y.Q., and H.B. designed the study; P.M.B., W.-H.L., S.B., M.R.R., and Y.Q. performed the experiments; and P.M.B., W.-H.L., Y.Q., and H.B. wrote the manuscript.

## Competing interests

H.B. is the founder of, a consultant for, and a shareholder of Oxford Nanopore Technologies, a company engaged in the development of nanopore sensing and sequencing technologies. P.M.B., W.-H.L., Y.Q., M.R.R., H.B., and S.G.-M., have filed patents describing the electroosmotically active nanopores and their applications of proteoform characterisation.

