## Supplemental information for "Enzyme-less nanopore detection of post-translational modifications within long polypeptides"

**Table S1 Sequences of the thioredoxin-linker concatemers**

| **Dimer (Trx-linker)$_2$** |
|---|
| CGSDKIIHLTDDSFDTDVLKADGAILVDFWAEWSGPSKMIAPILDEIADEYQGKLTVAKLNIDQNPGTAPKY<br>GIRGIPTLLLFKNGEVAATKVGALSKGQLKEFLDANLAGSAGSAGSAGSAGSAGSAGSAGSAGSAGRSDK<br>IIHLTDDSFDTDVLKADGAILVDFWAEWSGPSKMIAPILDEIADEYQGKLTVAKLNIDQNPGTAPKYGIRGIP<br>TLLLFKNGEVAATKVGALSKGQLKEFLDANLAGSAGSAGSAGSAGSAGSAGSAGSAGSAGRS |
| **Tetramer (Trx-linker)$_4$** |
| CGSDKIIHLTDDSFDTDVLKADGAILVDFWAEWSGPSKMIAPILDEIADEYQGKLTVAKLNIDQNPGTAPKY<br>GIRGIPTLLLFKNGEVAATKVGALSKGQLKEFLDANLAGSAGSAGSAGSAGSAGSAGSAGSAGSAGRSDK<br>IIHLTDDSFDTDVLKADGAILVDFWAEWSGPSKMIAPILDEIADEYQGKLTVAKLNIDQNPGTAPKYGIRGIP<br>TLLLFKNGEVAATKVGALSKGQLKEFLDANLAGSAGSAGSAGSAGSAGSAGSAGSAGSAGRSDKIIHLTD<br>DSFDTDVLKADGAILVDFWAEWSGPSKMIAPILDEIADEYQGKLTVAKLNIDQNPGTAPKYGIRGIPTLLLFK<br>NGEVAATKVGALSKGQLKEFLDANLAGSAGSAGSAGSAGSAGSAGSAGSAGSAGRSDKIIHLTDDSFDT<br>DVLKADGAILVDFWAEWSGPSKMIAPILDEIADEYQGKLTVAKLNIDQNPGTAPKYGIRGIPTLLLFKNGEVA<br>ATKVGALSKGQLKEFLDANLAGSAGSAGSAGSAGSAGSAGSAGSAGSAGRS |
| **Hexamer (Trx-linker)$_6$** |
| CGSDKIIHLTDDSFDTDVLKADGAILVDFWAEWSGPSKMIAPILDEIADEYQGKLTVAKLNIDQNPGTAPKY<br>GIRGIPTLLLFKNGEVAATKVGALSKGQLKEFLDANLAGSAGSAGSAGSAGSAGSAGSAGSAGSAGRSDK<br>IIHLTDDSFDTDVLKADGAILVDFWAEWSGPSKMIAPILDEIADEYQGKLTVAKLNIDQNPGTAPKYGIRGIP<br>TLLLFKNGEVAATKVGALSKGQLKEFLDANLAGSAGSAGSAGSAGSAGSAGSAGSAGSAGRSDKIIHLTD<br>DSFDTDVLKADGAILVDFWAEWSGPSKMIAPILDEIADEYQGKLTVAKLNIDQNPGTAPKYGIRGIPTLLLFK<br>NGEVAATKVGALSKGQLKEFLDANLAGSAGSAGSAGSAGSAGSAGSAGSAGSAGRSDKIIHLTDDSFDT<br>DVLKADGAILVDFWAEWSGPSKMIAPILDEIADEYQGKLTVAKLNIDQNPGTAPKYGIRGIPTLLLFKNGEVA<br>ATKVGALSKGQLKEFLDANLAGSAGSAGSAGSAGSAGSAGSAGSAGSAGRSDKIIHLTDDSFDTDVLKAD<br>GAILVDFWAEWSGPSKMIAPILDEIADEYQGKLTVAKLNIDQNPGTAPKYGIRGIPTLLLFKNGEVAATKVG<br>ALSKGQLKEFLDANLAGSAGSAGSAGSAGSAGSAGSAGSAGSAGRSDKIIHLTDDSFDTDVLKADGAILV<br>DFWAEWSGPSKMIAPILDEIADEYQGKLTVAKLNIDQNPGTAPKYGIRGIPTLLLFKNGEVAATKVGALSKG<br>QLKEFLDANLAGSAGSAGSAGSAGSAGSAGSAGSAGSAGRS |
| **Octamer (Trx-linker)$_8$** |
| CGSDKIIHLTDDSFDTDVLKADGAILVDFWAEWSGPSKMIAPILDEIADEYQGKLTVAKLNIDQNPGTAPKY<br>GIRGIPTLLLFKNGEVAATKVGALSKGQLKEFLDANLAGSAGSAGSAGSAGSAGSAGSAGSAGSAGRSDK<br>IIHLTDDSFDTDVLKADGAILVDFWAEWSGPSKMIAPILDEIADEYQGKLTVAKLNIDQNPGTAPKYGIRGIP<br>TLLLFKNGEVAATKVGALSKGQLKEFLDANLAGSAGSAGSAGSAGSAGSAGSAGSAGSAGRSDKIIHLTD<br>DSFDTDVLKADGAILVDFWAEWSGPSKMIAPILDEIADEYQGKLTVAKLNIDQNPGTAPKYGIRGIPTLLLFK<br>NGEVAATKVGALSKGQLKEFLDANLAGSAGSAGSAGSAGSAGSAGSAGSAGSAGRSDKIIHLTDDSFDT<br>DVLKADGAILVDFWAEWSGPSKMIAPILDEIADEYQGKLTVAKLNIDQNPGTAPKYGIRGIPTLLLFKNGEVA<br>ATKVGALSKGQLKEFLDANLAGSAGSAGSAGSAGSAGSAGSAGSAGSAGRSDKIIHLTDDSFDTDVLKAD<br>GAILVDFWAEWSGPSKMIAPILDEIADEYQGKLTVAKLNIDQNPGTAPKYGIRGIPTLLLFKNGEVAATKVG<br>ALSKGQLKEFLDANLAGSAGSAGSAGSAGSAGSAGSAGSAGSAGRSDKIIHLTDDSFDTDVLKADGAILV<br>DFWAEWSGPSKMIAPILDEIADEYQGKLTVAKLNIDQNPGTAPKYGIRGIPTLLLFKNGEVAATKVGALSKG<br>QLKEFLDANLAGSAGSAGSAGSAGSAGSAGSAGSAGSAGRSDKIIHLTDDSFDTDVLKADGAILVDFWAE<br>WSGPSKMIAPILDEIADEYQGKLTVAKLNIDQNPGTAPKYGIRGIPTLLLFKNGEVAATKVGALSKGQLKEFL<br>DANLAGSAGSAGSAGSAGSAGSAGSAGSAGSAGRSDKIIHLTDDSFDTDVLKADGAILVDFWAEWSGPS<br>KMIAPILDEIADEYQGKLTVAKLNIDQNPGTAPKYGIRGIPTLLLFKNGEVAATKVGALSKGQLKEFLDANLA |
| **Blue**: Trx; **Yellow**: linker; **Pink**: N-terminal cysteine-glycine; White: restriction enzyme site. |

| **Nonamer (Trx-linker)$_4$(Trx-linker-24S/26C)(Trx-linker)$_4$** |
|---|
| SDKIIHLTDDSFDTDVLKADGAILVDFWAEWSGPSKMIAPILDEIADEYQGKLTVAKLNIDQNPGTAPKYGIR<br>GIPTLLLFKNGEVAATKVGALSKGQLKEFLDANLAGSAGSAGSAGSAGSAGSAGSAGSAGSAGRSDKIIHL<br>TDDSFDTDVLKADGAILVDFWAEWSGPSKMIAPILDEIADEYQGKLTVAKLNIDQNPGTAPKYGIRGIPTLLL<br>FKNGEVAATKVGALSKGQLKEFLDANLAGSAGSAGSAGSAGSAGSAGSAGSAGSAGRSDKIIHLTDDSF<br>DTDVLKADGAILVDFWAEWSGPSKMIAPILDEIADEYQGKLTVAKLNIDQNPGTAPKYGIRGIPTLLLFKNGE<br>VAATKVGALSKGQLKEFLDANLAGSAGSAGSAGSAGSAGSAGSAGSAGSAGRSDKIIHLTDDSFDTDVLK<br>ADGAILVDFWAEWSGPSKMIAPILDEIADEYQGKLTVAKLNIDQNPGTAPKYGIRGIPTLLLFKNGEVAATKV<br>GALSKGQLKEFLDANLAGSAGSAGSAGSAGSAGSAGSAGSAGSAGRSGTSDKIIHLTDDSFDTDVLKAD<br>GAILVDFWAEWSGPSKMIAPILDEIADEYQGKLTVAKLNIDQNPGTAPKYGIRGIPTLLLFKNGEVAATKVG<br>ALSKGQLKEFLDANLAGSAGSAGSAGSAGSAGSAGS**RRASAC**AGSAGSAGRSPRRSDKIIHLTDDSFDT<br>DVLKADGAILVDFWAEWSGPSKMIAPILDEIADEYQGKLTVAKLNIDQNPGTAPKYGIRGIPTLLLFKNGEVA<br>ATKVGALSKGQLKEFLDANLAGSAGSAGSAGSAGSAGSAGSAGSAGSAGRSDKIIHLTDDSFDTDVLKAD<br>GAILVDFWAEWSGPSKMIAPILDEIADEYQGKLTVAKLNIDQNPGTAPKYGIRGIPTLLLFKNGEVAATKVG<br>ALSKGQLKEFLDANLAGSAGSAGSAGSAGSAGSAGSAGSAGSAGRSDKIIHLTDDSFDTDVLKADGAILV<br>DFWAEWSGPSKMIAPILDEIADEYQGKLTVAKLNIDQNPGTAPKYGIRGIPTLLLFKNGEVAATKVGALSKG<br>QLKEFLDANLAGSAGSAGSAGSAGSAGSAGSAGSAGSAGRSDKIIHLTDDSFDTDVLKADGAILVDFWAE<br>WSGPSKMIAPILDEIADEYQGKLTVAKLNIDQNPGTAPKYGIRGIPTLLLFKNGEVAATKVGALSKGQLKEFL<br>DANLAGSAGSAGSAGSAGSAGSAGSAGSAGSAGRS |
| **Nonamer (Trx-linker)$_4$(Trx-linker-14S/16C)(Trx-linker)$_4$** |
| SDKIIHLTDDSFDTDVLKADGAILVDFWAEWSGPSKMIAPILDEIADEYQGKLTVAKLNIDQNPGTAPKYGIR<br>GIPTLLLFKNGEVAATKVGALSKGQLKEFLDANLAGSAGSAGSAGSAGSAGSAGSAGSAGSAGRSDKIIHL<br>TDDSFDTDVLKADGAILVDFWAEWSGPSKMIAPILDEIADEYQGKLTVAKLNIDQNPGTAPKYGIRGIPTLLL<br>FKNGEVAATKVGALSKGQLKEFLDANLAGSAGSAGSAGSAGSAGSAGSAGSAGSAGRSDKIIHLTDDSF<br>DTDVLKADGAILVDFWAEWSGPSKMIAPILDEIADEYQGKLTVAKLNIDQNPGTAPKYGIRGIPTLLLFKNGE<br>VAATKVGALSKGQLKEFLDANLAGSAGSAGSAGSAGSAGSAGSAGSAGSAGRSDKIIHLTDDSFDTDVLK<br>ADGAILVDFWAEWSGPSKMIAPILDEIADEYQGKLTVAKLNIDQNPGTAPKYGIRGIPTLLLFKNGEVAATKV<br>GALSKGQLKEFLDANLAGSAGSAGSAGSAGSAGSAGSAGSAGSAGRSGTSDKIIHLTDDSFDTDVLKAD<br>GAILVDFWAEWSGPSKMIVPILDEIADEYQGKLTVAKLNIDQNPGTAPKYGIRGIPTLLLFKNGEVAATKVG<br>ALSKGQLKEFLDANLAGSAGSAGSAG**RRASAC**SAGSAGSAGSAGSAGSAGRSPRRSDKIIHLTDDSFDT<br>DVLKADGAILVDFWAEWSGPSKMIAPILDEIADEYQGKLTVAKLNIDQNPGTAPKYGIRGIPTLLLFKNGEVA<br>ATKVGALSKGQLKEFLDANLAGSAGSAGSAGSAGSAGSAGSAGSAGSAGRSDKIIHLTDDSFDTDVLKAD<br>GAILVDFWAEWSGPSKMIAPILDEIADEYQGKLTVAKLNIDQNPGTAPKYGIRGIPTLLLFKNGEVAATKVG<br>ALSKGQLKEFLDANLAGSAGSAGSAGSAGSAGSAGSAGSAGSAGRSDKIIHLTDDSFDTDVLKADGAILV<br>DFWAEWSGPSKMIAPILDEIADEYQGKLTVAKLNIDQNPGTAPKYGIRGIPTLLLFKNGEVAATKVGALSKG<br>QLKEFLDANLAGSAGSAGSAGSAGSAGSAGSAGSAGSAGRSDKIIHLTDDSFDTDVLKADGAILVDFWAE<br>WSGPSKMIAPILDEIADEYQGKLTVAKLNIDQNPGTAPKYGIRGIPTLLLFKNGEVAATKVGALSKGQLKEFL<br>DANLAGSAGSAGSAGSAGSAGSAGSAGSAGSAGRS |
| **Blue**: Trx; **Yellow**: linker; **Green**: modified linker; **Pink**: sequence for modification; White: restriction enzyme site. |

**Table S2 Percentage residual currents ($I_{res\%}$) for the three levels of repeating feature A recorded during C-terminus first concatemer translocation.**

| | Trx-linker dimer | Trx-linker tetramer | Trx-linker octamer |
|---|---|---|---|
| $I_{res\%}$ (A1)[a] | 34 ± 1% | 35 ± 1% | 35 ± 1% |
| $I_{res\%}$ (A2)[a] | 22 ± 3% | 24 ± 2% | 23 ± 1% |
| $I_{res\%}$ (A3)[a],[b] | 1.9 ± 2.4% | 1.7 ± 1.7% | 2.3 ± 2.7% |
| N[a],[b] | 105 units<br>2 separate pores | 66 units<br>3 separate pores | 443 units<br>2 separate pores |

**[a]** Ires% was calculated for each step in individual features A as the remaining current as a percentage of the open pore current (e.g., $I_{res\%}(A1) = I_{A1}/I_{open}$). The standard deviations were derived for N Trx-linker units collected with >1 separate pores. Conditions: 750 mM GdnHCl, 10 mM HEPES, pH 7.2, +140 mV (trans), 24 ± 1 °C.
**[b]** Trx-linker units that produced a Level A3 with a dwell time <1 ms were discarded during analysis. The associated spikey appearance suggested under-sampling and therefore an inaccurate $I_{res\%}$ value. Trx-linker units that generated a Level A3 with a dwell time >1 ms and a square shape were included in the $I_{res\%}$ analysis.

**Table S3 Frequency of C terminus-first or N terminus-first translocation events recorded with Trx-linker concatemers[a].**

| | Voltage (trans) | C terminus-first translocation | N terminus-first translocation | N |
|---|---|---|---|---|
| Dimer | +140 mV | 67% | 33% | 142 |
| Tetramer | +140 mV | 68% | 32% | 87 |
| Hexamer | +140 mV | 68% | 32% | 196 |
| Octamer | +120 mV | 86% | 14% | 87 |
| | +140 mV | 91% | 9% | 373 |
| | +160 mV | 94% | 6% | 192 |
| | +180 mV | 85% | 15% | 62 |

**[a]** Conditions: 750 mM GdnHCl, 10 mM HEPES, pH 7.2, the applied potential (trans) is specified in the table, 24 ± 1 °C. Concatemer concentrations: dimer 2.23 μM, tetramer 0.63 μM, hexamer 0.25 μM, octamer 0.81 μM.

**Table S4 Mean dwell times ($\langle\tau\rangle$) derived by QuB[a] for the three levels of repeating feature A (A1, A2, A3) recorded during the C-terminus first translocation of Trx-linker octamers through a single $(NN\_113R)_7$ nanopore[b]**

| Voltage (trans) | +140 mV | |
|---|---|---|
| **$\langle\tau_{A1}\rangle$ / ms** | 270 ± 20 | N = 277 |
| **$\langle\tau_{A2}\rangle$ / ms** | 23 ± 1 | N = 277 |
| **$\langle\tau_{A3}\rangle$ / ms** | 320 ± 60<br>0.69 ± 0.04 | N = 40<br>N = 294 |

**[a]** Dwell time analysis was performed by using the maximum interval likelihood algorithm of QuB.
**[b]** Conditions: 750 mM GdnHCl, 10 mM HEPES, pH 7.2, 24 ± 1 °C.

**Table S5 Percentage residual current ($I_{res\%}$) and root-mean-square noise ($I_{RMS}$) characteristics of individual modifications on Trx-linker nonamers.**

| | $\Delta I_{res\%}$[a] | $I_{RMS}$ / pA[c] | N |
|---|---|---|---|
| **Trx-linker-14S-P** | 4.4 ± 0.8% | 0.96 ± 0.18 | 19 concatemers<br>4 separate pores |
| | 4.0 ± 1.2%[b] | 1.6 ± 0.9[b] | 19 concatemers<br>4 separate pores |
| **Trx-linker-24S-P** | 8.3 ± 1.6% | 2.0 ± 0.6 | 27 concatemers<br>3 separate pores |
| | 9.2 ± 2.1%[b] | 2.5 ± 0.9[b] | 23 concatemers<br>3 separate pores |
| **Trx-linker-16C-GSH** | 5.1 ± 0.9% | 0.93 ± 0.19 | 46 concatemers<br>4 separate pores |
| **Trx-linker-26C-GSH** | 8.6 ± 1.3% | 1.6 ± 0.2 | 23 concatemers<br>3 separate pores |
| **Trx-linker-16C-SLN** | 15 ± 1% | 0.73 ± 0.28 | 24 concatemers<br>3 separate pores |
| **Trx-linker-26C-SLN** | 18 ± 2% | 1.8 ± 0.6 | 55 concatemers<br>5 separate pores |

**[a]** $\Delta I_{res\%}$ = <$I_{res\%}$(A1, Trx-linker)> – $I_{res\%}$(A1, Trx-linker+PTM). For a C terminus-first translocation event, <$I_{res\%}$(A1, Trx-linker)> was determined as the mean $I_{res\%}$ value of the unmodified A1 levels within an individual translocation event. $I_{res\%}$(A1, Trx-linker+PTM) was determined for the A1 level of the modified linker and appeared once per translocating concatamer. Conditions: 375 mM GdnHCl, 375 mM KCl, 10 mM HEPES, pH 7.2, +140 mV (trans), 24 ± 1 °C.
**[b]** Conditions: 750 mM GdnHCl, 10 mM HEPES, pH 7.2, +140 mV (trans), 24 ± 1 °C.
**[c]** Root-mean-square noise values ($I_{RMS}$) were measured from current traces after an applied post-recording filter at 2 kHz. $I_{RMS}$ was normalised by the noise of each pore ($I_{RMS}^2 = I_{RMS}$(A1, Trx-linker+PTM)$^2$ - $I_{RMS}$(open pore)$^2$).

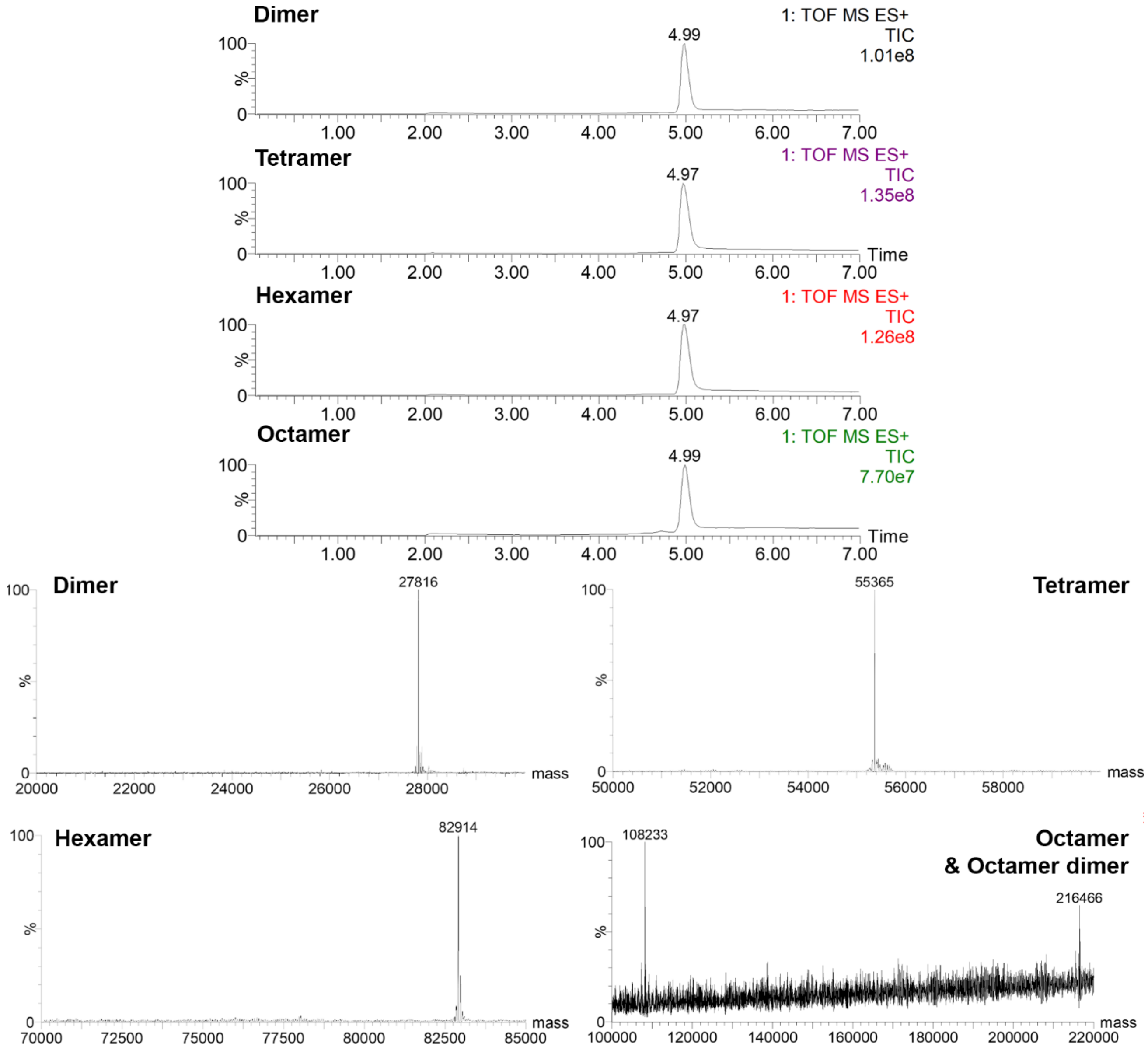


**Fig. S1 LC-MS characterization of Trx-linker concatemers**. LC-MS chromatograms (top) and deconvoluted ESI-MS spectra (bottom) are shown. Dimer, (Trx-linker)$_2$: mass = 27816 Da (calc) and 27816 Da (obs); Tetramer, (Trx-linker)$_4$: mass = 55367 Da (calc) and 55365 Da (obs); Hexamer, (Trx-linker)$_6$: mass = 82918 Da (calc) and 82914 Da (obs); Octamer, (Trx-linker)$_7$Trx: mass = 108231 Da (calc) and 108233 Da (obs); Dimer of octamers: mass = 216460 (calc) and 216466 Da (obs).

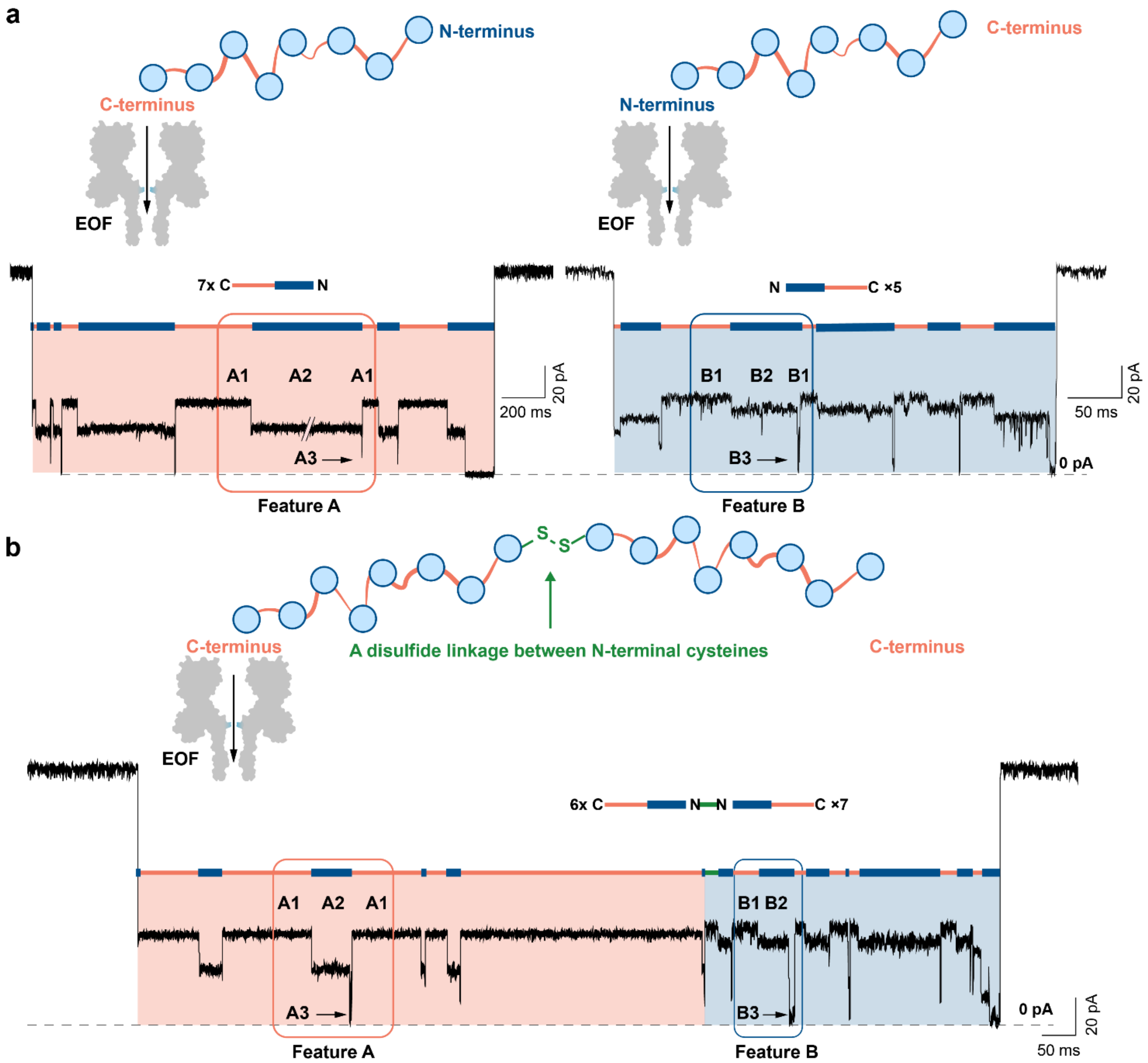


**Fig. S2 Repeating current features recorded during electroosmosis-driven concatemer translocation through a nanopore. a,** Two repeating current features, A or B, were recorded with Trx-linker octamers pre-treated with 5 mM tris(2-carboxyethyl)phosphine (TCEP) for 10 min prior to their addition to the cis compartment of the recording chamber. Conditions: 750 mM GdnHCl, 10 mM HEPES, pH 7.2, 0.81 μM Trx-linker octamer (cis), 5 mM TCEP, +140 mV (trans), 24 ± 1 °C. **b,** Without the TCEP pre-treatment, features A were always seen before features B when they occurred together within a single translocation event. The first two levels (B1 and B2) in features B have larger noise and higher $I_{res\%}$ compared to A1 and A2 recorded within a single translocation event with a single pore (A1: $I_{res\%}$= 35 ± 1 %, $I_{RMS}$ = 1.1 ± 0.1 pA, N = 25; A2: $I_{res\%}$ = 21 ± 1%, $I_{RMS}$ = 1.5 ± 0.2 pA, N = 25; B1: $I_{res\%}$= 38 ± 1 %, $I_{RMS}$ = 1.7 ± 0.4 pA, N = 39; B2: $I_{res\%}$ = 32 ± 1%, $I_{RMS}$ = 2.0 ± 0.5 pA, N = 39). The translocating molecules, which gave sequential A and B features, were assigned as dimers of octamers linked by a disulfide bond between the two N-terminal cysteines. Therefore, in the unlinked molecules (see '**a**'), C terminus-first translocation occurred when features A were observed and N terminus-first translocation occurred when features B were observed. The recorded repeating features are indicated by orange and blue bars. Conditions: 750 mM GdnHCl, 10 mM HEPES, pH 7.2, 0.81 μM Trx-linker octamer (cis), +140 mV (trans), 24 ± 1 °C. All traces were filtered at 2 kHz for clarity; transient A3 levels were truncated by filtering and therefore deviated from ~0 pA.

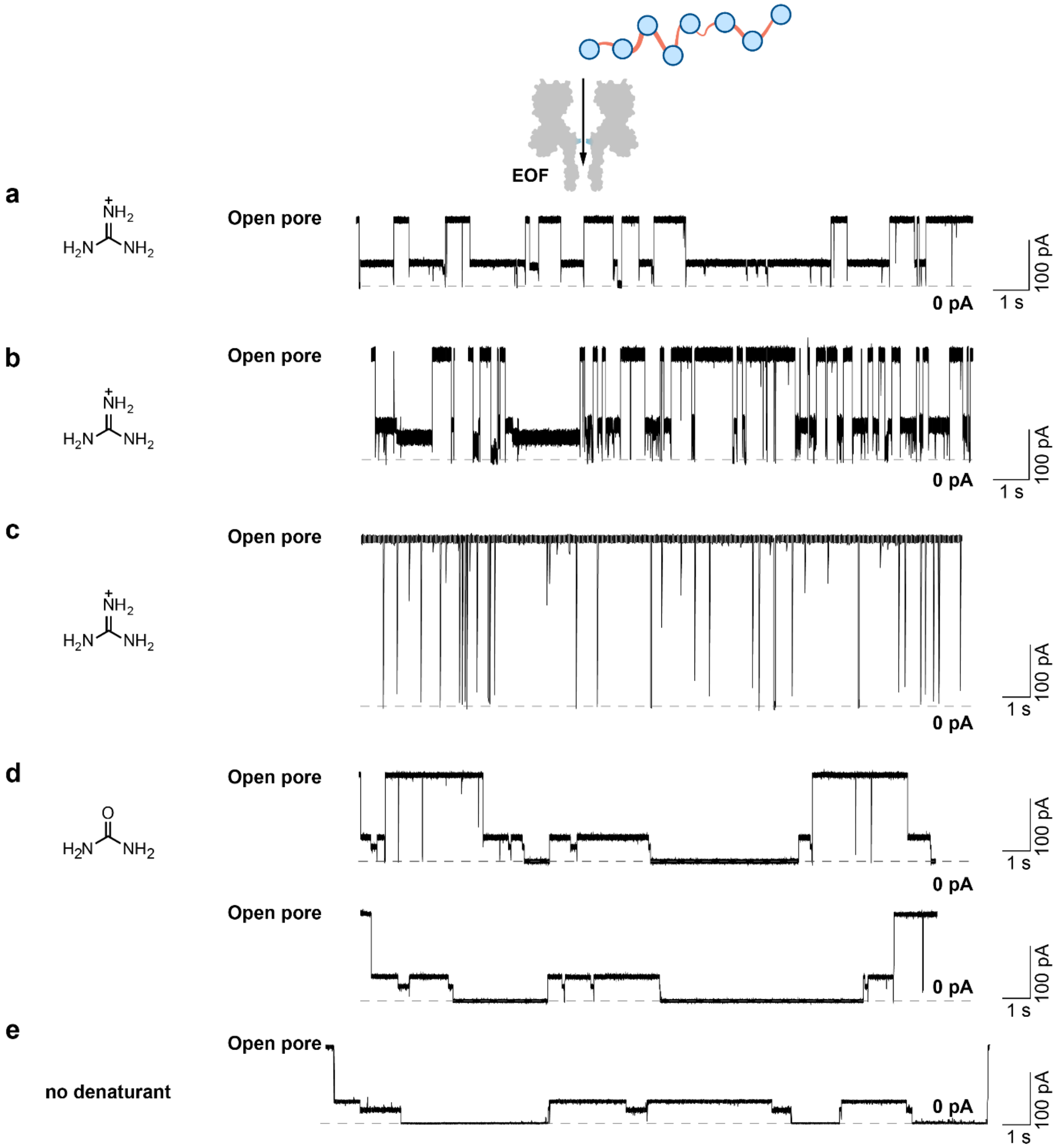


**Fig. S3 Electroosmosis-driven translocation of Trx-linker octamers through a nanopore. a-e,** Current traces for the translocation of Trx-linker octamers in the presence of 750 mM GdnHCl **(a)**, 1.5 M GdnHCl **(b)**, 3 M GdnHCl **(c)** without post-acquisition filtering, 2 M urea **(d)** or no denaturant **(e)** with 2 kHz post-acquisition filtering. Current features for subunit-by-subunit translocation were lost at 3 M GdnHCl **(c)**. The mean number of features A recorded per concatemer is **(a)** ~4, **(b)** ~3, **(c)** 0, **(d)** ~4, and **(e)** ~4. Conditions: 10 mM HEPES, pH 7.2, 0.81 μM Trx-linker octamer (cis), +140 mV (trans), 24 ± 1 °C, with **(a)** 750 mM GdnHCl; **(b)** 1.5 M GdnHCl; **(c)** 3 M GdnHCl; **(d)** 2 M urea and 750 mM KCl; **(e)** 750 mM KCl.

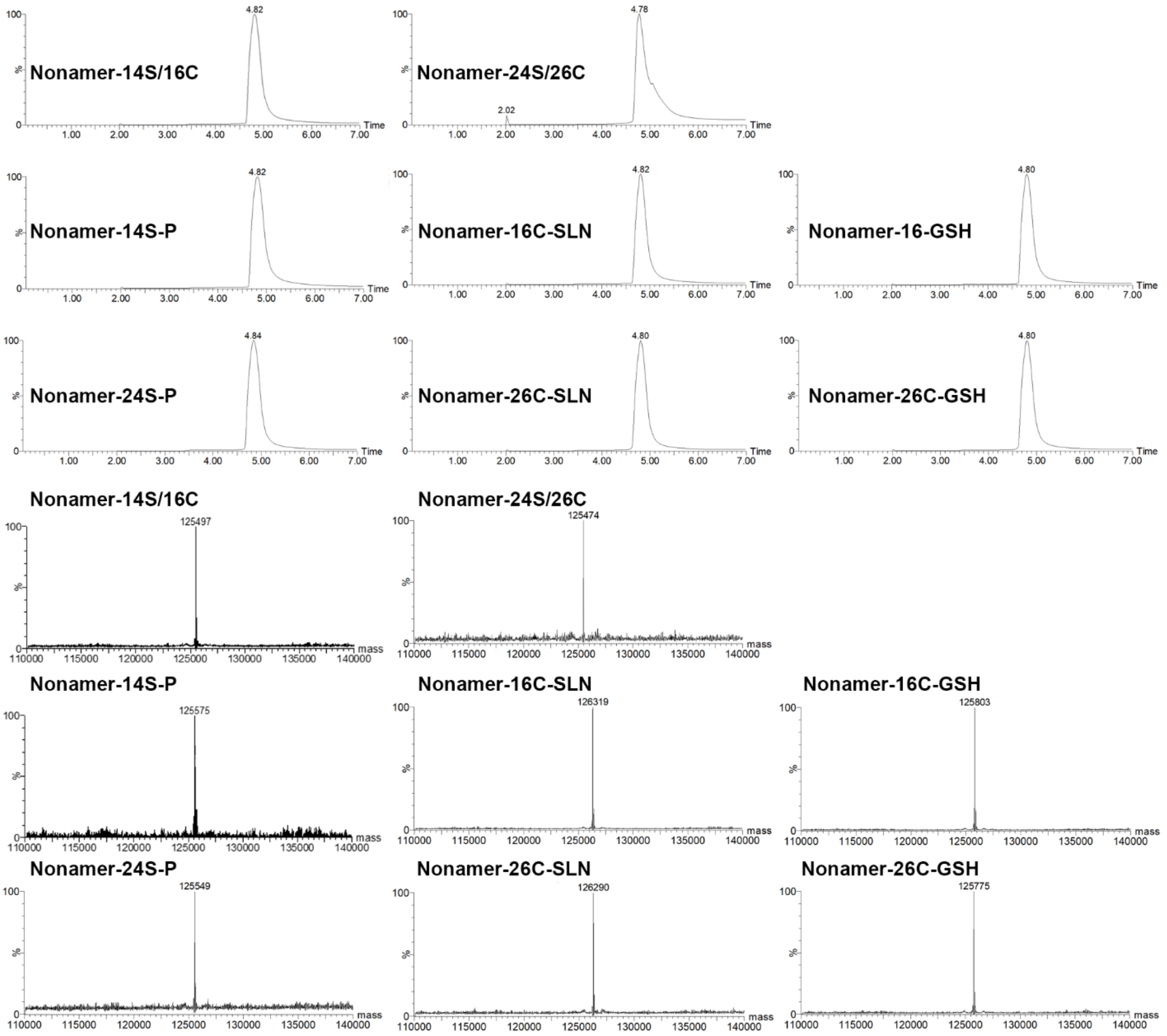


**Fig. S4 LC-MS characterization of Trx-linker nonamers**. LC-MS chromatograms (top) and deconvoluted ESI-MS spectra (bottom). Nonamer 14S/16C: mass = 125498 Da (calc) and 125497 Da (obs); Nonamer 24S/26C: mass = 125470 Da (calc) and 125474 Da (obs); Nonamer 14S-P: mass = 125578 Da (calc) and 125575 Da (obs); Nonamer 16C-SLN: mass = 126319 Da (calc) and 126319 Da (obs); Nonamer 16C-GSH: mass = 125803 (calc) and 125803 Da (obs); Nonamer 24S-P: mass = 125550 Da (calc) and 125549 Da (obs); Nonamer 26C-SLN: mass = 126291 Da (calc) and 126290 Da (obs); Nonamer 26C-GSH: mass = 125775 (calc) and 125775 Da (obs).

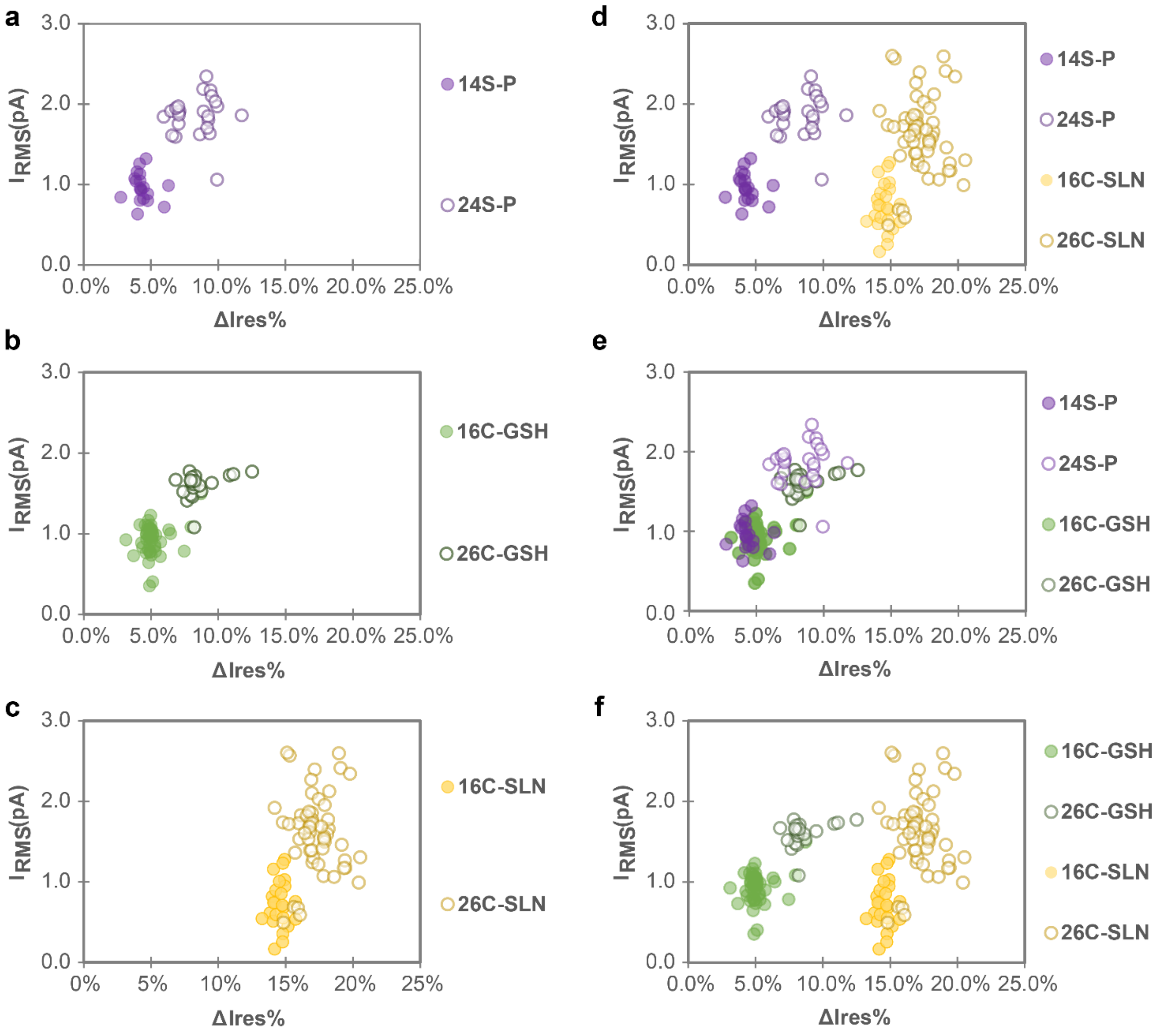


**Fig. S5 Identification and positional discrimination of PTMs in protein concatemers translocated by electroosmotic flow through a nanopore.** Protein nonamers containing a single PTM (See Fig. 3 for protein sequences and PTM structures) were tested. **a-c**, Scatter plots of $I_{RMS}$ and $\Delta I_{res\%}$ showing positional discrimination of a phosphorylated serine, a glutathionylated cysteine, or a glycosylated cysteine at sites 10 aa apart ($\Delta I_{res\%}$ = $<I_{res\%}$(A1, Trx-linker)$>$ – $I_{res\%}$(A1, Trx-linker+PTM), where $<I_{res\%}$(A1, Trx-linker)$>$ is the mean $I_{res\%}$ value of A1 levels of an unmodified unit within a single translocation event. Conditions: 375 mM GdnHCl, 375 mM KCl, 10 mM HEPES, pH 7.2, 1.2 µM Trx-linker nonamer (cis), +140 mV (trans), 24 ± 1 °C. **d-f,** Overlaid scatter plots of $I_{RMS}$ and $\Delta I_{res\%}$ showing discrimination between phosphorylated and glycosylated populations, glutathionylated and glycosylated populations, and overlaps between phosphorylated and glutathionylated populations.

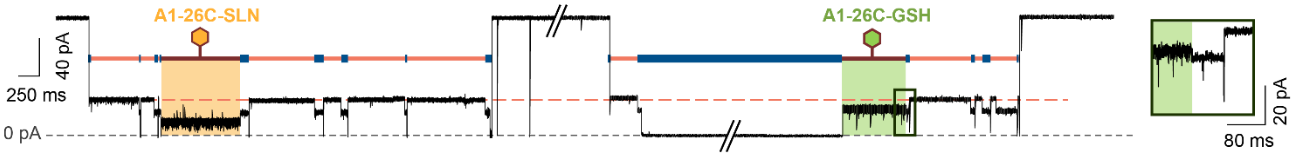


**Fig. S6 Identification of PTMs in a mixture of two concatemers.** Current traces recorded with the same nanopore for three C terminus-first translocations of a mixture of two Trx-linker nonamers containing either a GSH or SLN modification at position 26C within the central linker. The A1 level for the unmodified units (orange dash), and the A1 level for a unit modified with 26C-GSH (A1-26C-GSH, green) or 26-SLN (A1-26C-SLN, yellow) are colour-coded. The inset zooms in on the transition from A1-26C-GSH to A2. Traces have been filtered at 2 kHz. Conditions: 375 mM GdnHCl, 375 mM KCl, 10 mM HEPES, pH 7.2, 1.2 µM Trx-linker nonamer (cis), +140 mV (trans), 24 ± 1 °C.

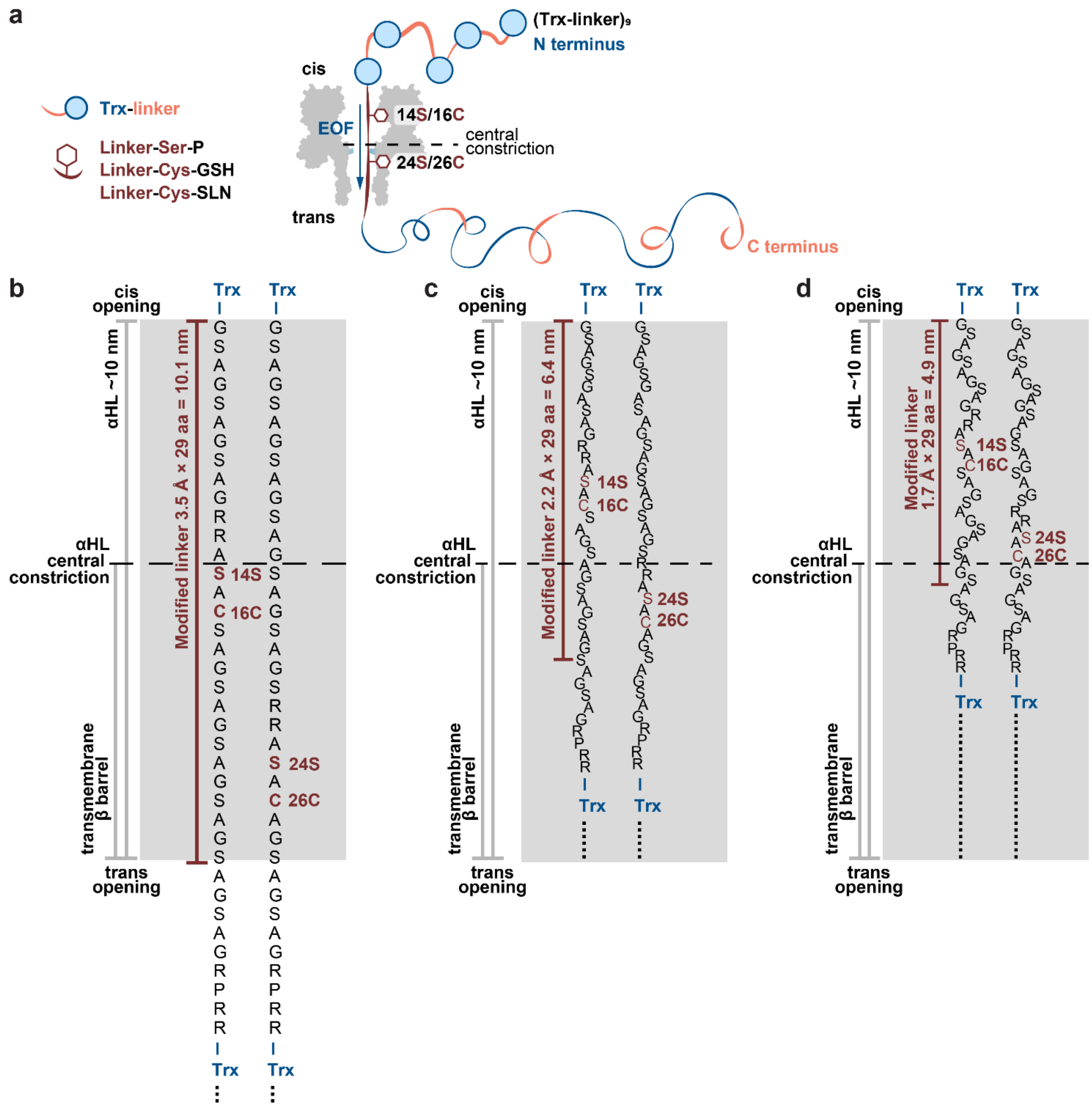


**Fig. S7 Positions of modification sites during translocation through an αHL pore. a,** The Trx-linker nonamers contained a RRASAC sequence within the central linker, which was post-translationally modified (hexagon). In a C-terminus-first threading configuration, as shown, the 14S/16C modification sites would be located closer to the cis opening of the αHL pore than the 24S/26C pair, when translocation is paused with a Trx unit at the cis mouth of the pore. **b-d,** Depending on the degree of extension of the polypeptide chain under the EOF (3.5 Å per aa when fully extended, 1.7-2.2 Å per aa under ~5-10 pN[1]), the 14S/16C and 24S/26C sites would be located at different positions within an αHL pore. Assuming that the N-terminal residue of the linker is at the cis opening of the pore when the translocation is arrested by a folded Trx unit, the modified linker (red) might fully span the αHL pore (**b**) or occupy only a part of the nanopore (**c**,**d**). When the 24S/26C sites are located nearer the central constriction of the αHL pore (**c**,**d**), a PTM at 24S/26C would produce a larger current blockade than that at 14S/16C (PTM = Ser-P, Cys-GSH, Cys-SLN), which is what is observed (Fig. 3b). Given that the applied potential drops mostly across the transmembrane β barrel[2], the current difference between 14S/16C+PTM and 24S/26C+PTM is likely to be larger in **c** than in **d**.
